# Small-scale field testing of Alpha-cypermethrin water-dispersible granules in comparison with the recommended wettable powder formulation for indoor residual spraying against malaria vectors in Benin

**DOI:** 10.1101/204651

**Authors:** Nicolas Moiroux, Armel Djenontin, Barnabas Zogo, Aziz Bouraima, Ibrahim Sidick, Olivier Pigeon, Cédric Pennetier

## Abstract

**Backgroud:** Pyrethroids are the most common class of insecticide used worldwide for indoor residual spraying (IRS) against malaria vectors. Water-dispersible granules (WG) are a pyrethroid formulation to be applied after disintegration and dispersion in water with less risks of inhalation than using the usual wettable powder (WP) formulation. The objective of this small-scale field study was to evaluate efficacy and duration of insecticidal action of a new alpha-cypermethrin WG (250g a.i./Kg) against susceptible *Anopheles gambiae* in comparison with the WHO reference product (alpha-cypermethrin WP, 50g a.i./Kg) on the most common indoor surfaces in Benin.

**Methods:** Both formulations were applied at two target-dose concentrations in houses made of mud and cement in the Tokoli village in southern Benin. We measured the applied dose of insecticide by chemical analysis of filter paper samples collected from the sprayed inner walls. We recorded *An. gambiae* mortality and knock-down rates every 15 days during 6 months using standard WHO bioassays.

**Results:** The alpha-cypermethrin WG formulation did not last as long as the WP formulation on both surfaces. The difference is higher with the 30mg/m^2^ concentration for which the WP formulation reached the 80% mortality threshold during 2 months on the mud-plastered walls (3 months on cement) whereas the WG formulation last only one month (2 months on cement).

**Conclusions:** The new WG formulation has a shorter efficacy than the WHO recommended WP formulation. In this trial, both the WG and WP formulations had low durations of efficacy that would need at least two rounds of spray to cover the entire transmission season.

## Background

During the last decade, Insecticide Treated Nets (ITN) became the major malaria vector control tool implemented in Africa, complemented by Indoor Residual Spraying (IRS) in some specific contexts. Indeed, these tools target different periods of the mosquito life cycle (host-seeking behavior and resting behavior, respectively). in 2015, National Malaria control programs (NMCP) reported that about 106 million people worldwide were protected by IRS. Pyrethroids were the class of insecticides the most used for IRS [1]. Over the 59 countries that have implemented IRS in 2014, 43 declared using pyrethroids alone or in combination with other classes of insecticides [2].

Pyrethroids insecticides are usually available in wettable powder (WP) formulations that present some disadvantages. First, the particles in suspensions made from wettable powders are large and visible residues may be left on sprayed surfaces. Moreover, there is a risk of inhalation during mixing, as the dry particles can become airborne. Alternatively, water-dispersible granules (WG) are a formulation consisting of granules to be applied by spraying after disintegration and dispersion in water. There is less risk of inhalation of airborne particles from water-dispersible granules than from wettable powders [3].

Alpha-cypermethrin is among the 12 insecticides recommended by the World Health Organization Pesticide Scheme (WHOPES) for IRS [4]. Alpha-cypermethrin has been tested and recommended by the World Health Organization (WHO) as a wettable powder (WP) and aqueous suspension concentrate at a dosage between 20 and 30mg/m^2^ with expected residual activity of 4 to 6 months [5].

A WG formulation of alpha-cypermethrin was tested in india [6]. This formulation showed residual efficacy for 13–15 weeks for the 20 mg/m^2^ application rate and 13–16 weeks for the 30 mg/m^2^ rate not significantly different than the WHO recommended alpha-cypermethrin WP formulation [7].

The objective of the present small-scale field study was to evaluate efficacy and duration of insecticidal action of a new alpha-cypermethrin WG (250g a.i./Kg) at two application rates (20 and 30 mg ai/m^2^) against susceptible *Anopheles gambiae* in comparison with similar dosages of the reference product (alpha-cypermethrin WP, 50g a.i./Kg) on the most common indoor surfaces in Benin.

## Methods

### Study area

The study was carried out in Tokoli (6°27’30” N; 2°10’16” E) in the Ouidah-Kpomasse-Tori health district in southern Benin. The climate is sub-equatorial with two dry seasons (August-September and December-March) and two rainy seasons (April-July and October-November). The average annual rainfall is around 1200 mm. The average monthly temperatures vary between 27 and 31°C.

Houses in southern Benin are of three types [8] that are all present in Tokoli: Mud-plastered houses, houses made with white sand and cement, and houses made with red sand and cement.

### Insecticides tested

α-cypermethrin WG: wettable granule formulation of alpha-cypermethrin (250 g a.i./kg) manufactured by Tagros Chemicals India Limited, Chennai, Jhaver Centre, Rajah Annamalai Bulding, IV Floor, 72, Marshalls Road, Egmore Chennai-600 008,India (Batch number AG-01/13).

α-cypermethrin WP: WHOPES recommended alpha-cypermethrin Wettable Powder (50g a.i./Kg).

### Study design

As previous findings showed no difference in insecticide efficacy and persistence applied on red or white sands mixed with cement [8], we retained only two types of house (mud-plastered and cement) for this study. We included in the study 50 houses (25 mud-plastered and 25 with cement walls) of 112 present in the village. In each batch of 25, houses were randomly allocated to one of the 5 following arms (i.e. 5 houses per arm):

- α-cypermethrin WG at the 20 mg/m^2^ ±25% target dose (WG20)
- α-cypermethrin WG at the 30 mg/m^2^ ±25% target dose (WG30)
- α-cypermethrin WP at the 20 mg/m^2^ ±25% target dose (WP20)
- α-cypermethrin WP at the 30 mg/m^2^ ±25% target dose (WP30)
- control (not sprayed)

Houses were sprayed between the 7 and 16 march 2014 at the end of the long dry season. Only one room per house was sprayed. insecticide was applied once, using a hand-operated compression sprayer equipped with a flat fan nozzle 80° swath and 0.76 L/min flow rate. The four walls were treated. The spraying was done by two well trained technicians. They attended a 4-days training with a WHOPES mandated expert just before the beginning of spraying.

### Safety precautions and ethical considerations

The National Research Ethics Committee of Benin approved the study (reference number 016 of July 16th, 2013). Householders and spray men gave their informed consent.

Safety precautions regarding mixing, handling and spraying the insecticide followed standard WHOPES procedures as outlined in [9]. Spray men used recommended/necessary protective clothing. They were given an information sheet in French and were briefed on possible adverse effects and the need to fully comply with safety instructions. Spray men were advised that in the event of any discomfort, they would be subjected to medical examination and care.

Rooms were sprayed after the personal effects of the householders have been removed and/or protected by craft paper. The windows, the floor, and the doors were also protected by craft paper during the spray.

The householders were advised about safety precautions in order to avoid any risks during and after the spray. They were advised to remain out of the rooms during the spray and up to 3 hours after spraying. They were told that it is required to protect them from coming in contact with fumes of the insecticide spray. The adult householders were advised to ask their children not to intentionally touch the sprayed walls for at least one day after spraying since the walls remained wet for about a day. After a room has been sprayed, it is essential that walls are not scrubbed or mutilated or plastered until the end of the study. The householders were therefore advised not to do so as part of the informed consent form. The householders were also advised that in the event of an adverse effect or illness due to fever, they should approach the Medical Officer at the closest health centre for treatment but may also seek advice/assistance from our institutions (IRD and CREC) at the contact details given in the consent form.

### Adverse effects on spray men and householders

Spray men were interviewed using a questionnaire at the end of a day of spraying, the following morning and one week after. Moreover, our team visited each household one week and one month after spraying to record adverse effect on the inhabitants using a questionnaire administered to the household heads.

### Residual activity

Standard WHO bioassays [10] were carried out on days 1, 15, 30, 45, 60, 90, 120, 150 and 180 after spraying, using laboratory-reared, susceptible females of *An. gambiae* (Kisumu strain). Batches of 10 unfed mosquitoes, 3–5 days old were put in a standard WHO cone and applied for 30 minutes (exposure time) on the four walls and the roof of each selected room. After 60 minutes, knock down (KD) mosquitoes were counted and all mosquitoes were kept for 24 hours in the laboratory to assess mortality at 24 hours.

### Chemical analysis

Before spraying, Whatman papers of 10 x 10 cm size were attached to the four inner walls of each of the selected rooms before spraying, and collected 24h post-spraying for chemical residue analysis. Positions of filter paper on the wall were marked to avoid carrying out cone bioassays at such spots.

Each paper sample was packed in aluminum foil separately and put in labeled bags. The packed samples was stored in a fridge at +4 ^0^C temperature before sending them to the WHO collaborating centre, Gembloux, Belgium for α-cypermethrin dosage.

### Statistical analysis

The α-cypermethrin content measured by chemical analysis on filter papers was compared between the 20mg/m^2^ and 30mg/m^2^ spraying objectives (for both formulations and both surfaces) and between the WP and WG formulations (for both spraying objectives and both surfaces) using T-tests.

Mortality and KD rates measured with WHO bioassays were analysed using binomial regression models. Due to convergence issue, we failed to fit mixed effect models with a random intercept for houses (to deal with possible auto-correlation among bioassay performed in a same room). We therefore fitted classical binomial-response regression models with the treatment, the surface (mud or cement), the time after spraying (log transformed) and interactions as fixed effects but without random intercept. The ‘brglm’ function of the ‘brglm’ package [11] in the software ‘R’ [12] was used for this analysis. It allows to fit binomial-response regression models using the bias-reduction method developed by Firth [13]. These procedures return estimates with improved frequentist properties (bias, mean squared error) that are always finite even in cases where the maximum likelihood estimates are infinite (data separation). The fitted models were used to predict (using the ‘predict’ function in R) the day when mortality fall significantly under the efficacy threshold of 80%.

## Results

On cement walls, chemical analysis of filter papers indicates that the mean concentrations of WP20 and WP30 were 32.60 mg/m^2^ (95% CI [27.39 - 37.80]) and 43.92 mg/m^2^ (95% CI [36.62 - 51.21])’ respectively (Figure 1A). On the same surface, the mean concentrations of WG20 and WG30 were 38.45 mg/m^2^ (95% CI [31.48 - 45.41]) and 53.47 mg/m^2^ (95% CI [44.53 - 62.42]), respectively (Figure 1C). The differences in α-cypermethrin contents between the WP and WG formulations applied on cement walls were not significant (p = 0.167 and p = 0.091 for the 20 and 30 mg/m^2^ target doses, respectively). On mud walls, chemical analysis of filter papers indicates that the mean concentration of WP20 and WP30 were 36.62 mg/m^2^ (95% CI [30.91 - 42.32]) and 41.77 mg/m^2^ (95% CI [35.55 - 47.99])’ respectively (Figure IB). On the same surface, the mean concentration of WG20 and WG30 were 37.23 mg/m^2^ (95% CI [30.31 - 44.14]) and 39.79 mg/m^2^ (95% CI [33.47 - 46.11]), respectively (Figure 1D). The differences in α-cypermethrin contents between the WP and WG formulations applied on mud walls were not significant (p = 0.961 and p = 0.559 for the 20 and 30 mg/m^2^ target doses, respectively). The mean applied to target dose ratio was 1.64 (95% CI [1.61 - 1.67]), exceeding expectation. As expected, α-cypermethrin contents on cement wall were significantly higher with the 30mg/m^2^ targeted doses (for both the WG and WP formulations) than with the 20mg/m^2^ target dose (Figures 1A and 1C). However on mud, we were not able to find any difference between papers from rooms sprayed at 20mg/m^2^ or 30 mg/m^2^ (Figures 1B and 1D).

**Figure 1:**
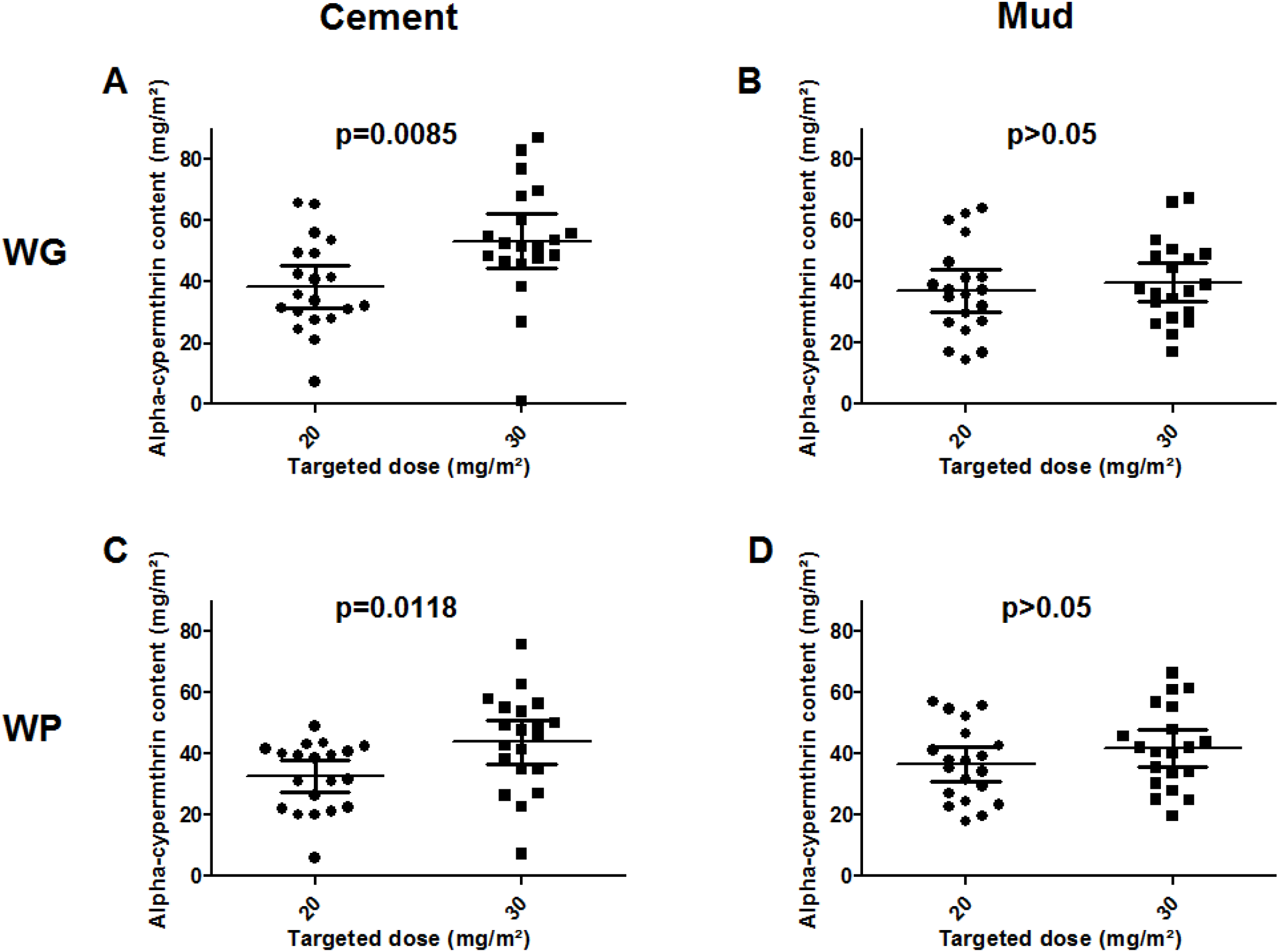
Comparison of applied and target doses (20 or 30 mg/m^2^) of insecticide according to the formulation (WP or WG) and the wall surface (mud or ciment). Mean with 95% confidence interval are shown. T-test p-value are provided.

Over 40 filter papers coming from the 10 control houses, 39 showed a concentration of alpha-cypermethrin lower than the limit of detection (i.e. < 4mg/m^2^) and one was just above (6.9 mg/m^2^).

On the mud plastered walls, the mortality model indicated that the WG20 and WP20 treatments efficacy failed significantly under the 80% threshold 26 and 27 days after spraying, respectively (Figure 2A). The WG30 treatment was efficient until the 30^th^ day after spraying. In contrast, the reference WP30 treatment was significantly more persistent with an induced mortality ≥80% until the 60^st^ day (Figure2C). The same trends were observed for the KD rate (Fig. 3A and 3C) except for the WP30 formulation that maintained a KD rate ≥ 80% until the 80^th^ day.

**Figure 2:**
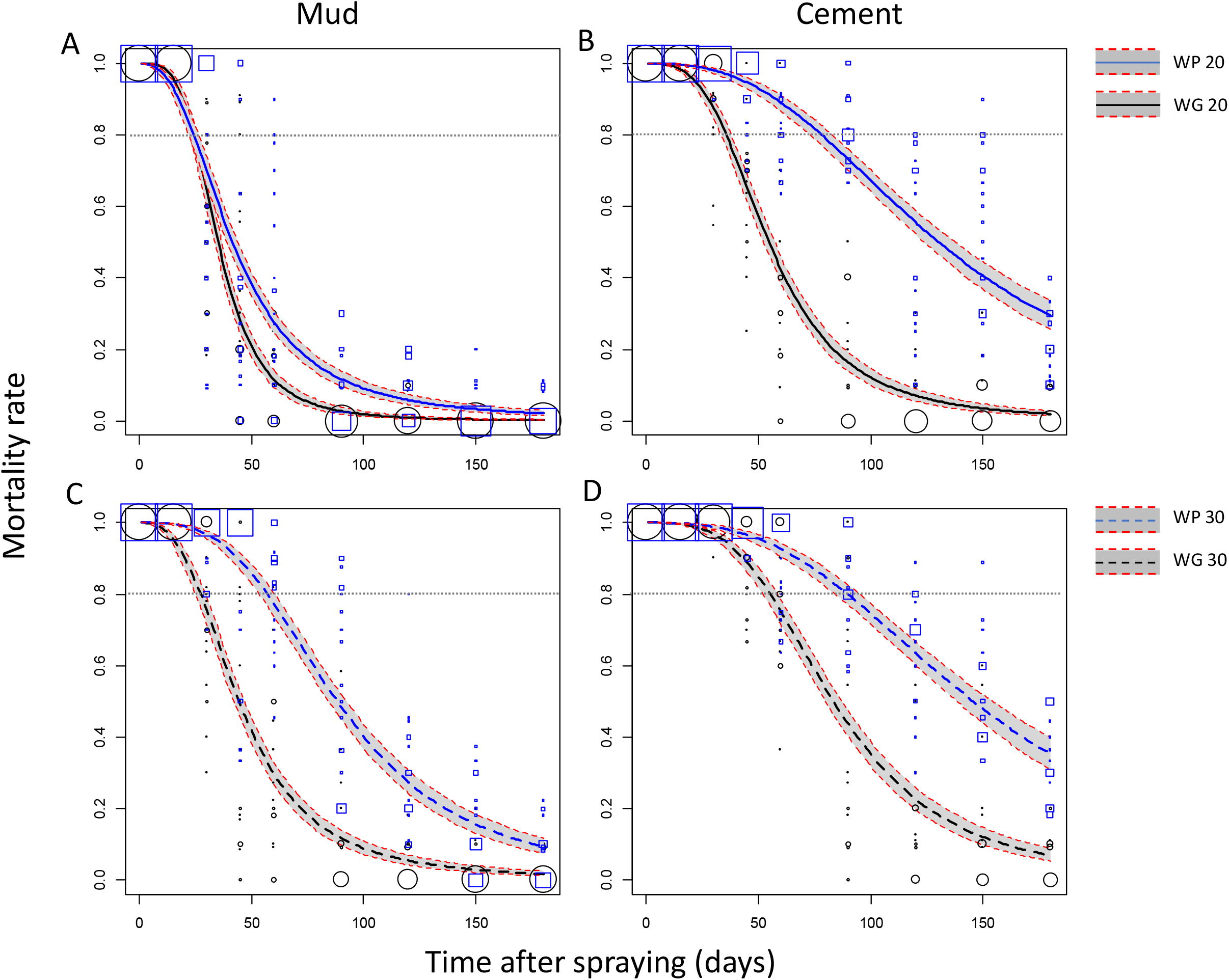
Efficacy (mortality) over time of Indoor Residual Spraying of the WP and WG formulations of Alphacypermethrin against susceptible *An. gambiae*. Mortality rates were predicted from a binomial-response regression model. Formulation WP (blue lines) and WG (black lines) at the 20 (A, B; solid lines) or 30 mg/m^2^ (C, D; dashed lines) targeted dose applied on mud (A, C) or cement walls (B, D) are compared. Grey areas are 95% confidence interval of predicted means. Mortality values measured on the field and used to fit the regression model are shown as blue squares (WP) and black circles (WG) of size proportional to the number of values (max = 20).

**Figure 3:**
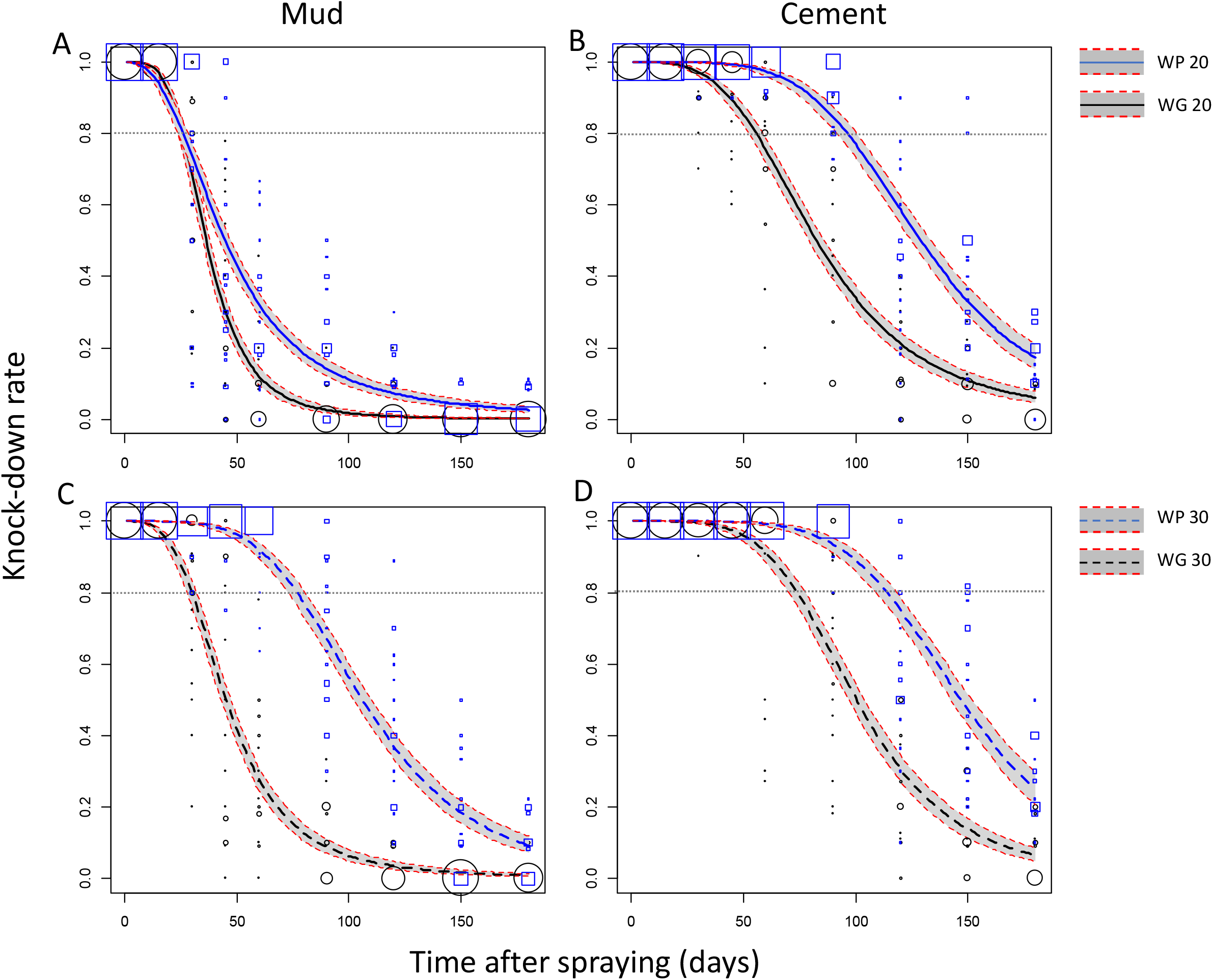
Efficacy (knock-down effect) over time of Indoor Residual Spraying of the WP and WG formulations of Alphacypermethrin against susceptible *An. gambiae*. Knock-down (KD) rates were predicted from a binomial-response regression model. Formulation WP (blue lines) and WG (black lines) at the 20 (A, B; solid lines) or 30 mg/m^2^ (C, D; dashed lines) targeted dose applied on mud (A, C) or cement walls (B, D) are compared. Grey areas are 95% confidence interval of predicted means. KD values measured on the field and used to fit the regression model are shown as blue squares (WP) and black circles (WG) of size proportional to the number of values (max = 20).

On the cement walls, mortalities induced by the WG formulation failed under the 80% threshold after 38 and 59 days when applied at the 20 mg/m^2^ and 30 mg/m^2^ target doses, respectively (Figure 2B and 2D). In comparison, mortalities induced by the reference WP formulation failed under the 80% threshold after 83 and 96 days when applied at 20 and 30 mg/m^2^, respectively (Fig. 2B and 2D). Regarding the KD rate induced by the WG formulation, it failed under 80% after 59 and 78 days when applied at the 20 and 30 mg/m^2^, respectively (Fig. 3B and 3D). With the reference WP formulation, the KD rate failed under 80% after 101 and 119 days for the 20 and 30 mg/m^2^ target doses, respectively (Fig. 3C and 3D). In the control rooms, the mean mortality rate was 0.0077 (95% CI [0.005 – 0.01]) 24h post exposure.

Two weeks after spraying, 6 household heads over 40 receiving an IRS treatment declared having experienced adverse effects (Table 1). They declared having experienced skin itching (N=2), runny nose (N=3), sneezing (N=3), eye watering (N=1), headache (N=2) and nausea-vomiting-stomach pain (N=1). One month after spraying, only two household heads having received the WG30 or WP30 treatments declared having experienced adverse effects (runny nose (N=1), sneezing (N=2), eye watering (N=1)). Spray men did not reported any adverse effect as well as householder of the control arm.

**Table 1:**
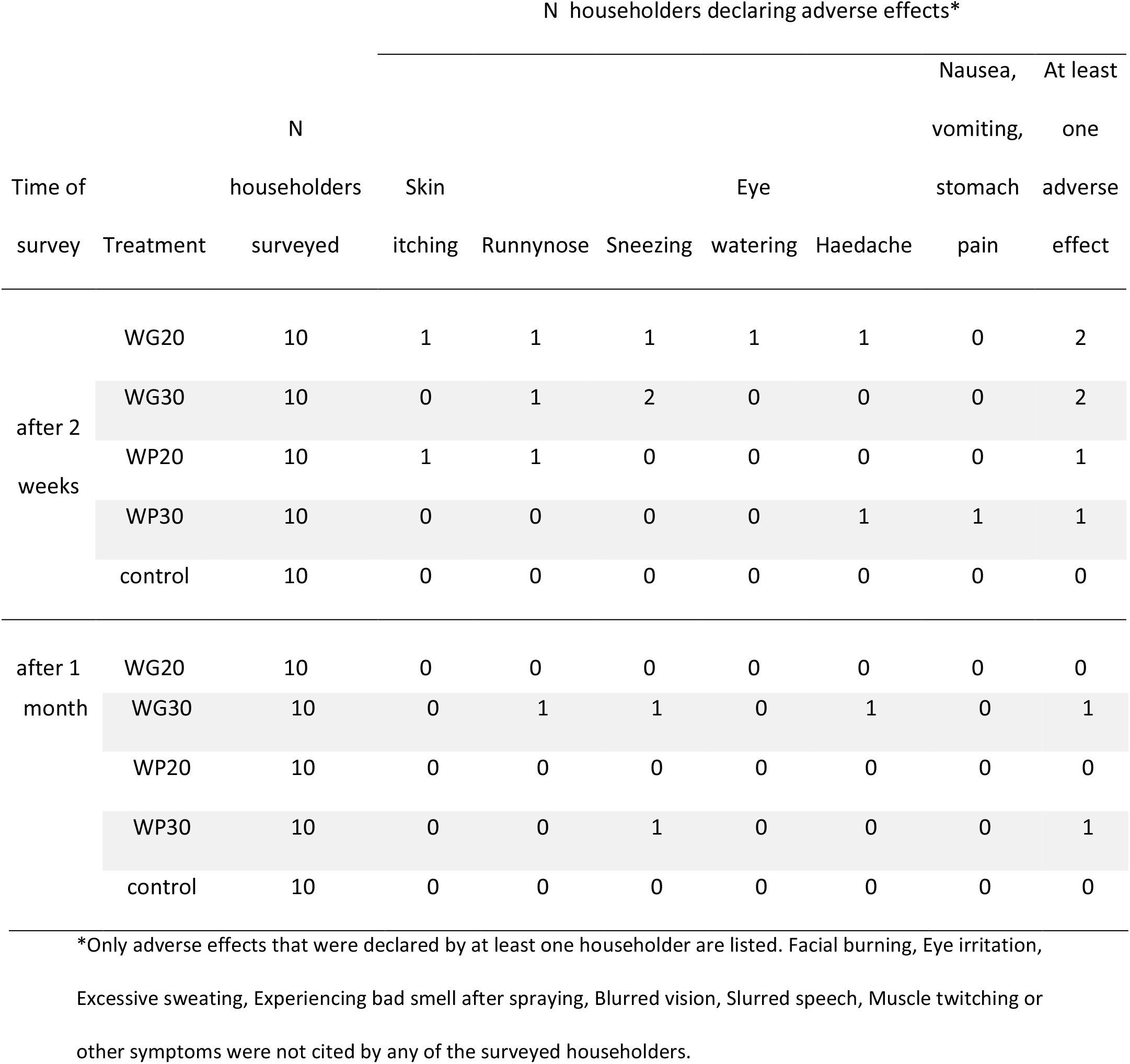
Number of householders declaring adverse effects two weeks and one month after spraying according to the treatment arm.

## Discussion

In the present study, the residual efficacy of a new WG formulation of α-cypermethrin for IRS was tested on most common indoor surfaces (mud and cement) against a susceptible strain of *Anopheles gambiae* in Benin and was compared to the WHO recommended WP formulation.

This small-scale trial showed that the α-cypermethrin WG formulation efficacy did not last as long as the WP formulation on both surfaces. The difference is higher with the 30mg/m^2^ concentration for which the WP formulation reached the 80% mortality threshold during 2 months whereas the WG formulation last only one month on the mud-plastered walls. The same trend is observed for the cement surface on which the efficacy was ≥ 80% mortality during approximately 3 months for the WP formulation and less than 2 months for the WG formulation. This indicate that the new WG formulation was not as persistent as the reference WP formulation. These results contrasts with those of Uragayala *et al*. [6] in india who found that the WG formulation was as efficient as the reference WP formulation with durations of efficacy higher than 3 months whatever the target dose or the wall surface. Such differences observed between Benin and india might be due to the different Anopheles species used for bioassay, differences in wall surfaces and in climatic conditions (the indian trial was performed during the dry season while the Beninese trial was almost entirely performed during the rainy season).

The short residual efficacy of both formulation in our trial indicated that multiple rounds of spraying should be needed to protect population for the entire malaria season that everywhere exceed 4 months in Benin. Because IRS are expensive to implement for resource-limited countries [14], such interventions should be limited to areas where the transmission season is short (e.g. in the sahelian area). Moreover, we found a difference in residual efficacy between the mud and cement surfaces. This observation, that was already made in southern Benin [15], highlights the potential difference in residual efficacy of IRS between rural areas where the majority of houses is mud-plastered and the urban areas where cement surfaces are more common.

In tropical environments, mosquitoes that are KD have little chances to recover because of the biomass of potential predators and scavengers (ants, spies, geckos…) [16]. The KD rate is therefore a highly relevant criteria for evaluation of IRS residual efficacy.

The chemical analysis showed a high variability of α-cypermethrin content within the filter papers attached on the inner walls, despite the great experience of the sprayers and the training they attended just before the sprays (with a WHOPES mandated expert). In operational conditions of IRS implementation when local operators are less experienced, we can therefore expect high variability in insecticide concentration applied in the houses. Independent evaluation of spraying quality should help improve operational procedures and go with Phase III evaluation studies [10].

It is notable that almost all α-cypermethrin contents measured on filter papers were above the targeted doses. However, the contents measured in houses that received the WP20 and WG20 treatment where still in the WHOPES recommended application dose of 20–30 mg/m^2^ ± 25% [7]. This makes the efficacy results reliable.

On cement wall, we find a positive relationship between the targeted dose and the concentrations measured on filter papers. It was expected to find the same on mud walls but we were not able to evidence such a relationship. We therefore hypothesise that an unknown proportion of the insecticide migrates into the mud wall by adsorption or absorption. if confirmed, this issue might be easily solved by inserting an inert plastic sheet between the wall and the filter paper.

Adverse effects were reported only in houses having received an IRS treatment. However, sample sizes (i.e. 10 households per arm) were two small to allow statistical comparisons between arms with a sufficient power.

## Conclusion

The tested α-cypermethrinWG formulation applied at a WHOPES recommended dose of 20–30 mg/m^2^ ± 25% (i.e. the WG20 and WP20 treatments) reached the cut-off point of 80% mortality during less than 2 months whatever the wall surface. This efficacy level was lesser than the WHO recommended α-cypermethrin WP formulation (almost 3 months before failing under 80% mortality) when applied on cement walls. When applied on mud-plastered walls, both formulations failed to exceed one month of efficacy. Because of these low durations of efficacy, we do not recommend the use of these formulations in Benin where more than 2 rounds of spray should be needed to cover the entire transmission season.

## Declarations

### Ethics approval and consent to participate

The National Research Ethics Committee of Benin approved the study (reference number 016 of July l6th, 2013). Householders and spray men gave their informed consent.

### Consent for publication

Not Applicable

### Availability of data and material

The datasets generated and/or analysed during the current study are not publicly available. The present study was performed under Technical Service Agreements (TSA) between iRD and WHO. According to these TSAs, IRD should not disclose the data without the authorization of WHO. Readers may contact WHOPES office to request the data at the following email address

### Authors’ contributions

CP and AD designed the study. BZ, AB and IS participated in data collection and performed WHO bioassay. OP performed and criticized chemical analysis. NM analyzed the data. NM, AD and CP interpreted the data. NM and AD drafted the manuscript. All authors read and approved the final manuscript.

## Acknowledgements

We thank populations of the study area, Tokoli Village in the Ouidah-Kpomasse-Tori health district, for their collaboration.

## List of abbreviations

IRD: institut de Recherche pour le Développement
CREC: Centre de Recherche en Entomologie de Cotonou
IRS: indoor Residual Spraying
WG: Water-dispersible Granules
WP: Wettable Powder
WHO: World Health Organisation
WHOPES: World Health Organisation Pesticides Evaluation Schema
NMCP: National Malaria Control Program
ITN: insecticide Treated Net
a.i.: active ingredient
KD: Knock Down
CI: Confidence interval

## References

1. World Health Organization. World malaria report 2016. Geneva: World Health Organization; 2016. http://www.who.int/iris/handle/10665/252038. Accessed 8 Sep 2017.

2. World Health Organization. World malaria report 2015. Geneva: World Health Organization; 2015. http://www.who.int/iris/handle/10665/200018. Accessed 6 Sep 2016.

3. WHO. Pesticides and their application: for the control of vectors and pests of public health importance. Geneva: World Health Organization; 2006.

4. World Health Organization. WHO recommended insecticides for indoor residual spraying against malaria vectors. 2016. http://www.who.int/whopes/Insecticides_IRS_2_March_2015.pd. Accessed 15 Sep 2016.

5. WHO Pesticide Evaluation Scheme. Report of the second WHOPES working group meeting. Geneva: World Health Organization; 1998. http://www.who.int/iris/handle/10665/64438. Accessed 15 Sep 2016.

6. Uragayala S, Kamaraju R, Tiwari S, Ghosh SK, Valecha N. Small-scale evaluation of the efficacy and residual activity of alpha-cypermethrin WG (250 g AI/kg) for indoor spraying in comparison with alpha-cypermethrin WP (50 g AI/kg) in India. Malar J. 2015;14:223.

7. WHO Pesticide Evaluation Scheme. Report of the seventeenth WHOPES working group meeting. Geneva: World Health Organization; 2014. http://www.who.int/iris/handle/10665/137514. Accessed 15 Sep 2016.

8. Djenontin A. Stratégies de gestion de la résistance aux insecticides des vecteurs du paludisme et impact opérationnel en Afrique de l’Ouest. Université d’Abomey-Calavi; 2011.

9. World Health Organization. Indoor residual spraying: an operational manual for indoor residual spraying (IRS) for malaria transmission control and elimination. 2nd edition. Geneva: World Health Organization; 2015. http://www.who.int/iris/handle/10665/177242. Accessed 15 Sep 2016.

10. WHO. Guidelines for testing mosquito adulticides intended for Indoor Residual Spraying (IRS) and Insecticide Treated Nets (ITNs). 2006;WHO/CDS/NTD/WHOPES/GCDDP/2006.3.

11. Kosmidis I. brglm: Bias reduction in binomial-response Generalized Linear Models. 2013. http://www.ucl.ac.uk/~ucakiko/software.html.

12. R Development Core Team. R: A Language and Environment for Statistical Computing. Vienna, Austria: R Foundation for Statistical Computing; 2010. http://www.R-project.org.

13. Firth D. Bias reduction of maximum likelihood estimates. Biometrika. 1993;80:20–38.

14. White MT, Conteh L, Cibulskis R, Ghani AC. Costs and cost-effectiveness of malaria control interventions - a systematic review. Malar J. 2011;10:1–14.

15. Djènontin A, Aïmihouè O, Sèzonlin M, Damien GB, Ossè R, Soukou B, et al. The residual life of bendiocarb on different substrates under laboratory and field conditions in Benin, Western Africa. BMC Res Notes. 2013;6:458.

16. Najera JA, Zaim M, WHO CPE Dept., WHOPES. Malaria vector control: insecticides for indoor residual spraying. Geneva: World Health Organization; 2001. http://www.who.int/iris/handle/l0665/67383. Accessed 15 Sep 2016.

